# Rotational Activity Around an Obstacle in 2D Cardiac Tissue in Presence of Cellular Heterogeneity

**DOI:** 10.1101/2021.09.28.462113

**Authors:** Pavel Konovalov, Daria Mangileva, Arsenii Dokuchaev, Olga Solovyova, Alexander V. Panfilov

## Abstract

Waves of electrical excitation rotating around an obstacle is one of the important mechanisms of dangerous cardiac arrhythmias occurring in the heart damaged by post-infarction scar. Such a scar also has a border zone around it, which has electrophysiological properties different from the rest of normal myocardial tissue. Spatial patterns of wave rotation in the presence of such tissue heterogeneity are poorly studied. In this paper we perform a comprehensive numerical study of various regimes of rotation of a wave in a plane layer of the ventricular tissue around an obstacle surrounded by a gray zone. We use a TP06 cellular ionic model which reproduces the electrophysiological properties of cardiomyocytes in the left ventricle of human heart. We vary the extent of obstacle and gray zone and study the pattern of wave rotation and its period. We observed different regimes of wave rotation that can be subdivided into several classes: (1) functional rotation and (2) scar rotation regimes, which were identified in the previous studies, and new (3) gray zone rotation regime: where the wave instead of rotation around the obstacle, rotates around the gray zone (an area of tissue heterogeneity) itself. For each class, the period of rotation is determined by different factors, which we discuss and quantify. We also found that due to regional pathological remodeling of myocardial tissue, we can obtain additional regimes associated with dynamical instabilities of two types which may affect or not affect the period of rotation.

## 1. Introduction

Propagating non-linear waves occur in various types of excitable media of physical [1], chemical [2,3] and biological nature[4,5]. These waves can also form vortices which organize spatial excitation patterns in the medium and their onset can have important consequences. For example, rotating waves in the heart is the main mechanism of dangerous cardiac arrhythmias [6,7] and study such regimes in the heart is important area of research in cardiac electrophysiology. In some patients, arrhythmias may arise from myocardial infarction, i.e. condition when due to poor blood supply a region of the heart is damaged. As a result, electrical waves propagating through the heart break and form vortices rotating in/around the damaged area [8,9].

From mathematical point such regimes can be viewed as rotation of non-linear waves around an obstacle. This is one of the classical regimes studied in the theory of excitable media. Such regimes were described in cellular automata models as early as in 1946 [10]. Later they were studied in reaction-diffusion models of excitable media [11]. More recently interest in such regimes, especially on rotation around obstacle of small size was initiated by the concept of anchoring of vortices proposed by Davidenko and co-authors [12]. Such processes were also studied using theoretical approaches [13,14].

Unfortunately, direct application of such studies to arrhythmias which occur around infarcted area is difficult, due to the following problem. Post-infarction scar area in the myocardial tissue has a complex geometry and functional heterogeneity [15]. It contains a compact scar region, which is similar to unexcitable obstacle. Such compact scar is surrounded by a border zone, which has both cellular and tissue properties different from the rest of normal cardiac tissue [16]. This zone is also called gray zone (GZ) because of its intermediate signal intensity in clinical MRI [17]. In mathematical sense this problem can be viewed as wave rotation around an obstacle surrounded by a heterogeneous media. There is a study by Bernus and co-authors [18] on the heterogeneity in the gray zone which triggers such rotating wave. But this study focused on initiation of arrhythmia, and it does not describe regimes of wave rotation after initiation. Such rotation regimes are poorly understood.

Main aim of this paper is to perform a generic numerical study of this problem. In this way, we use a simplified 2D representation of ventricular tissue with an obstacle and gray zone of simple circular geometry, which gives us only two parameters (obstacle and gray zone radii) to vary the extent of injury. In particular, we study rotation of waves in such 2D models with unexcitable obstacles surrounded by a border region where the ionic properties of cardiomyocytes are different form those in the rest of the tissue. We vary the size of the obstacle and gray zone to reveal their effects on the pattern of wave rotation and its period, which is an important characteristic of cardiac arrhythmia. We show that depending on the parameters, the wave can rotate either around the compact scar, or other regimes occur where the wave rotates within the gray zone or on its border with the normal tissue. The switching between these regimes can be understood from the principle of minimal period described in this paper. The dependency of the period on the size of the scar in the physiologically relevant range is determined by the slope of the conduction velocity restitution curve and can be estimated from a simple equation derived in the paper. We also show that for small scars the regional tissue heterogeneity can induce dynamical instability. Overall we provide classification of excitation patterns and determine main factors which can affect the period of the wave rotation.

## 2. Materials and Methods

### 2.1. Baseline ventricular myocardial tissue model. Numerical approach and software

To describe propagation of the excitation wave in the ventricular myocardial tissue from human heart, we used a 3-dimensional monodomain approach [19] and isotropic medium:

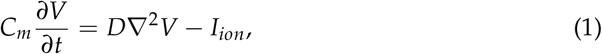

where *V* is the transmembrane potential; D is electro-diffusion matrix. Here, we used a uniform D value throughout the tissue simulating isotropic media. *C_m_* is the capacitance of the cell membrane. *I_ion_* is sum of all transmembrane ionic currents, described with biophysically detailed cellular ionic TP06 model of human ventricular cardiomyocytes [20]:

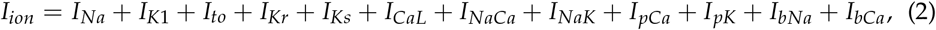

where *I_Na_* is the *Na*^+^ current, *I*_*K*1_ is the inward rectifier *K*^+^ current, *I_to_* is the transient outward current, *I_Kr_* is the delayed rectifier current, *I_Ks_* is the slow delayed rectifier current, *I_CaL_* is the L-type *Ca*^2+^ current, *I_NaK_* is the *Na*^+^/*Ca*^2+^ exchanger current, *I_NaK_* is the *Na*^+^/*K*^+^ ATPhase current, *I_pCa_* and *I_pK_* are plateau *Ca*^2+^ and *K*^+^ currents, and *I_bNa_* and *I_bCa_* are background *Na*^+^ and *Ca*^2+^ currents. This cellular model provides a detailed description of voltage, ionic currents, and intracellular ion concentrations and is based on a wide range of human electrophysiological data. The properties of all these currents are fitted to their experimentally measured values and their dynamics is fitted using additional differential equations. Each of the currents typically depends on a maximal constant conductivity *G_max_*, and on current values of time-dependent *V* and certain gating variables.

In this study we used a thin single layer of excitable tissue simulating 2D myocardial slab. Simulations were performed on grids 500 × 500 × 1 elements (140 × 140 × 0, 28 mm^3^) or 750 × 750 × 1 elements (210*mm* × 210 × 0, 28 mm^3^). Each tissue element can represent either excitable myocardial tissue or inexcitable post-infaction scar. Larger grids were used for modelling post-infarction scars with radius *r_IS_* > 16.8 mm.

Boundary conditions were formulated as the no flux through the boundaries:

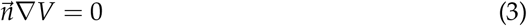

where 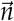 is the normal to the boundary. Infarction scar elements were simulated as non-conducting inexcitable obstacles and considered as internal boundaries (no flux) for the myocardial elements.

### 2.2. Simulation of post-infarction scar and gray zone

Post-infarction scar (IS) was modelled as a non-conducting inexcitable circular obstacle at the center of the tissue slab (black area in Fig. 1, left). It is surrounded by a gray zone (gray area in Fig. 1, left). Here, we used a TP06 cellular ionic model [20] with referent parameters for every cell in the healthy myocardial tissue, and a model with modified parameters (Table 1) for every cell in the gray zone reflecting cellular remodeling in myocardial tissue around the scar. Modification of TP06 ionic model for description of cells in the gray zone was taken from [21] and is similar to the properties of gray zone cells from [22]. In particular, the maximal conductances of several ionic currents are changed against the reference values as specified in Table 1. The difference in model parameters result in the difference in the action potential shape and duration between the cells from gray zone and healthy myocardium (Fig. 1(right)). The action potential duration is longer in the gray zone. The difference in the cellular activity between the border zone and the rest of the healthy tissue provides the functional heterogeneity in myocardial tissue around the scar.

**Figure 1.**
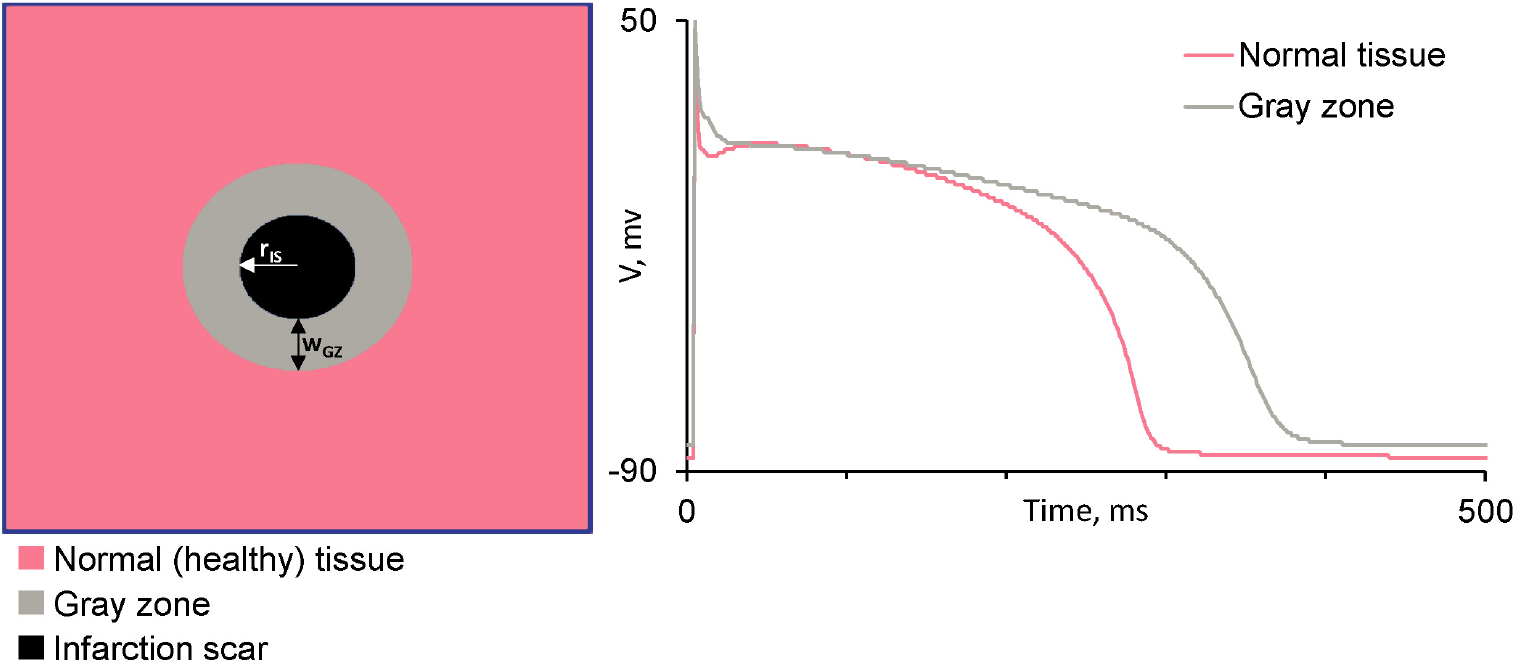
Left: Schematic representation of the geometry of cardiac tissue with the post-infarction scar: Normal tissue (pink), post-infarction scar (black) and gray zone around it (gray). Right: the shape of the action potential vs time in cells of healthy myocardium (pink) and gray zone (gray).

**Table 1:**
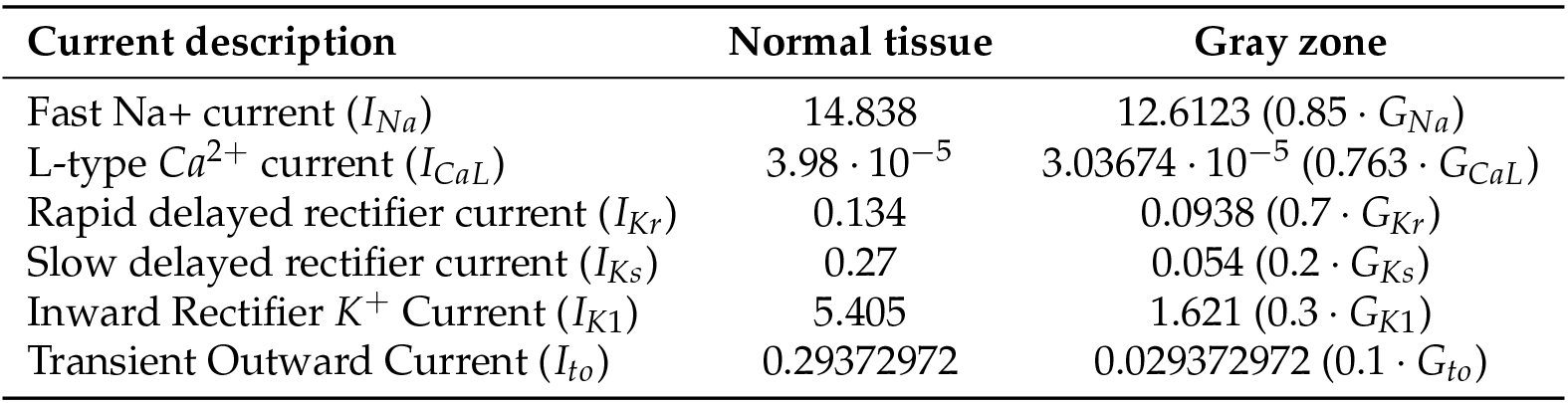
Maximal conductivity (*G*_*_) of ionic currents in cellular models from the normal tissue (referent values) and gray zone (modified values).

| Current description | Normal tissue | Gray zone |
| --- | --- | --- |
| Fast Na<sup>+</sup> current ($I_{Na}$) | 14.838 | 12.6123 ($0.85 \cdot G_{Na}$) |
| L-type Ca<sup>2+</sup> current ($I_{CaL}$) | $3.98 \cdot 10^{-5}$ | $3.03674 \cdot 10^{-5}$ ($0.763 \cdot G_{CaL}$) |
| Rapid delayed rectifier current ($I_{Kr}$) | 0.134 | 0.0938 ($0.7 \cdot G_{Kr}$) |
| Slow delayed rectifier current ($I_{Ks}$) | 0.27 | 0.054 ($0.2 \cdot G_{Ks}$) |
| Inward Rectifier K<sup>+</sup> Current ($I_{K1}$) | 5.405 | 1.621 ($0.3 \cdot G_{K1}$) |
| Transient Outward Current ($I_{to}$) | 0.29372972 | 0.029372972 ($0.1 \cdot G_{to}$) |

### 2.3. Protocol of numerical experiments in 2D tissue

A rotating wave in a 2D myocardial tissue slab with a post-infarction scar and gray zone was initiated counterclockwise using a standard protocol S1S2 (see Fig. S1 in the Supplement for graphical description of the protocol). Period of the stabilized wave was calculated in an arbitrary point in the healthy tissue as a time interval between the subsequent moments of the membrane depolarisation (zero *V* value at V^′^ *>*0).

We varied the size of scar from *r_IS_* = 0 to *r_IS_* = 53.2 mm with a step of 2.8 mm, and width of the gray zone from *w_GZ_* = 0 to *w_GZ_* = 42 mm also with a step of 2.8 mm and analysed the wave rotation pattern and evaluated the rotation period.

Note, in this study, we did not simulate the variability in the cellular population within the tissue. Here, we assumed two uniform areas of the myocardial tissue around the scar with the reference cellular model used throughout the healthy myocardium and the modified cellular model used throughout the gray zone. Thus, a single model run per each fixed pair (*r_IS_*, *w_GZ_*) of varied infarction parameters was performed.

#### Numerical methods

To solve the problem (1)-(3) we used a finite-difference method with 9-point stencil discretization scheme as described in [23] with 0.28 mm for the spatial step and 0.02 ms for the time step. Rush-Larsen formalism [24] was used for TP06 gating variables integration.

All calculations were performed on a C program on clusters “URAN” (IMM, Ural Branch of RAS) and “IIP” of Institute of immunology and Physiology (Ekaterinburg). The program uses CUDA for GPU parallelization and was compiled with a Nvidia C Compiler “nvcc”. Computational nodes have graphical cards Tesla K40m0.

## 3 Results

### 3.1. Wave rotation regimes

Varying the radius of post-infarction scar (*r_IS_*) and gray zone width (*w_GZ_*) in our 2D tissue models, we observed several qualitatively different spatial patterns of the wave rotation. Representative types of the wave patterns are shown in Figure 2.

**Figure 2.**
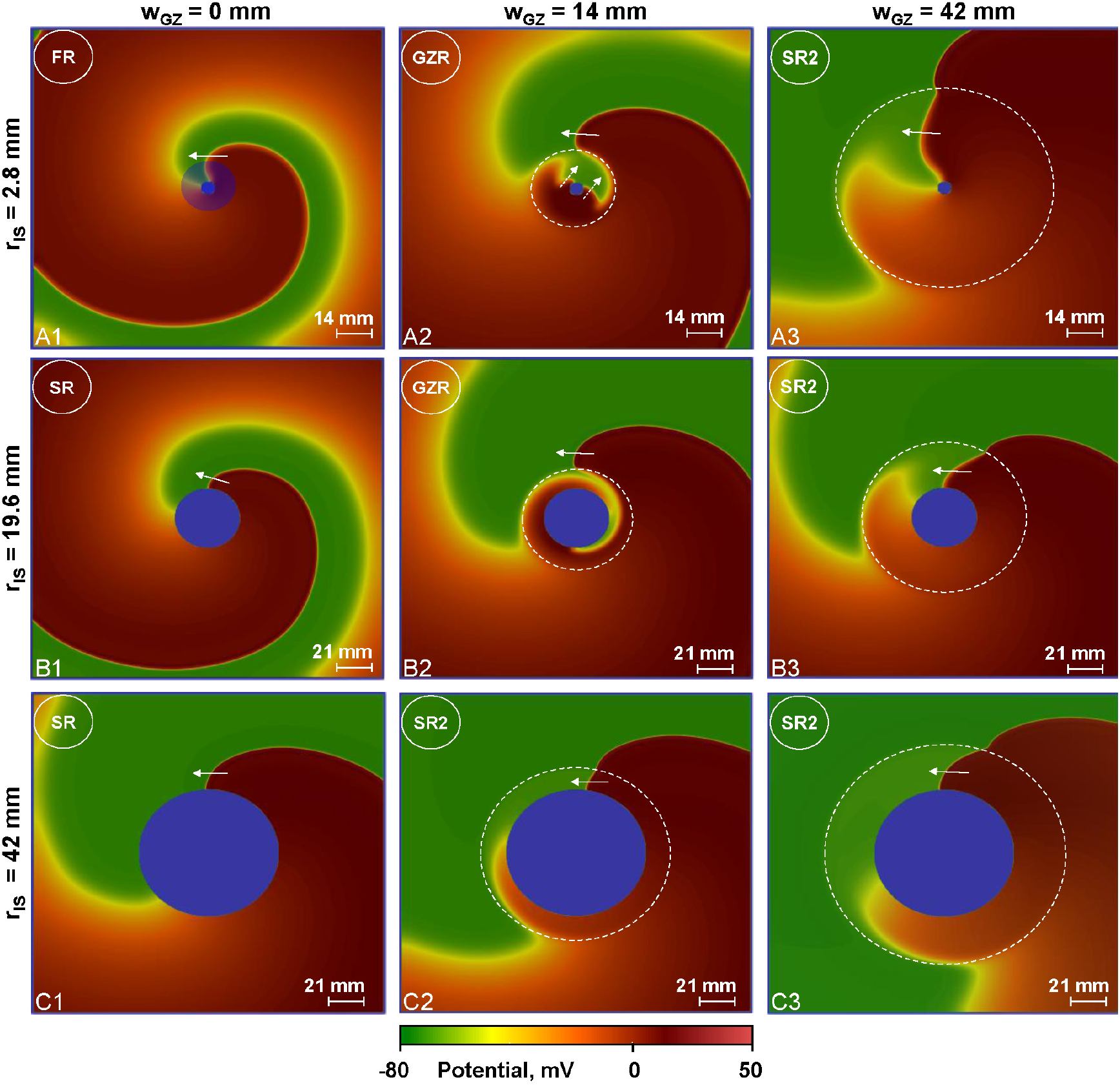
Examples of waves rotating around post-infarction scars of different sizes (up to down: *r_IS_* = 2.8*mm*; 19.6*mm*; 42*mm*) with a gray zone of different sizes (left to right: *w_GZ_* = 0*mm*; 14*mm*; 42*mm*). Four rotation regimes are shown: *scar rotation* (SR), *functional rotation* (FR, blue shading shows the effective obstacle around which the wave rotates), *gray zone rotation* (GZR), and *scar rotation 2* (SR2). See text for the description of the rotation regimes. Left column: typical view of the wave rotating around the scar in the absence of gray zone (*w_GZ_* = 0 mm). Waves rotate in two regimes: *FR* (A1) and *SR* (B1, C1). Middle column: waves rotate around the gray zone in a *GZR* (A2, B2) and *SR2* (C2). Right column: (*w_GZ_* = 42 mm) *SR2* regime in all cases (A3-C3). Arrows show the direction of the wave rotation. Dashed arrows in A2 (*r_IS_* = 19, 6*mm*, *w_GZ_* = 14*mm*) show wave direction inside the gray zone. Dashed circles show the border of the gray zones. Movies of rotation of spiral waves in *GZR* and *SR2* regimes can be found in the Supplement (movies S1 and S2).

In the absence of gray zone (Fig. 2 B1, C1, *w_GZ_* = 0 mm) the rotation occurs with the wavefront orthogonal to the boundary of the scar. Let us call this regime as a *scar rotation*, as here the radius or rotation is determined by the radius of the scar. However, for a small scar in Figure 2.A1 the situation is different. It looks as if the wave is rotating around an effective obstacle of a larger radius: the point where displacement in the normal direction coincides with the visible rotation around a center is located on a circle of a larger radius than the scar (blue shaded circle). Let us call such regime as a *functional rotation*. That means that the radius of rotation here is determined by the wave itself and not by the obstacle.

For the gray zone of *w_GZ_* = 14 mm (Fig.2 A2, B2) the rotation occurs with the wavefront orthogonal to the boundary of the gray zone, as if it is non-excitable. Inside the gray zone we see an area in red-orange colors indicating that spatial duration of the action potential here in prolonged. Let us call this regime as a *gray zone rotation* as here the radius or rotation is determined by the radius of the gray zone. For the largest scar in Fig.2(C2) the wavefront is attached and almost orthogonal to the scar and we observe scar rotation, however, as the wave here moves inside the gray zone let us call this regime as a *scar rotation 2*.

For the largest gray zones of *w_GZ_* = 42 mm (Fig.2 B3, C3) we see the *scar rotation 2* regime, except that in Fig.2(A3), where we see some features of *functional rotation* inside the gray zone.

Now let us analyse the period of rotation, which is one of the main characteristics of cardiac arrhythmia.

### 3.2. Rotation period depending on geometry of the scar

We plotted dependencies of the rotation period on the radius of the scar for 4 values of the gray zone width (Fig. 3) and dependencies of the rotation period on the width of the gray zone for 4 values of the scar radius (Fig. 4).

**Figure 3.**
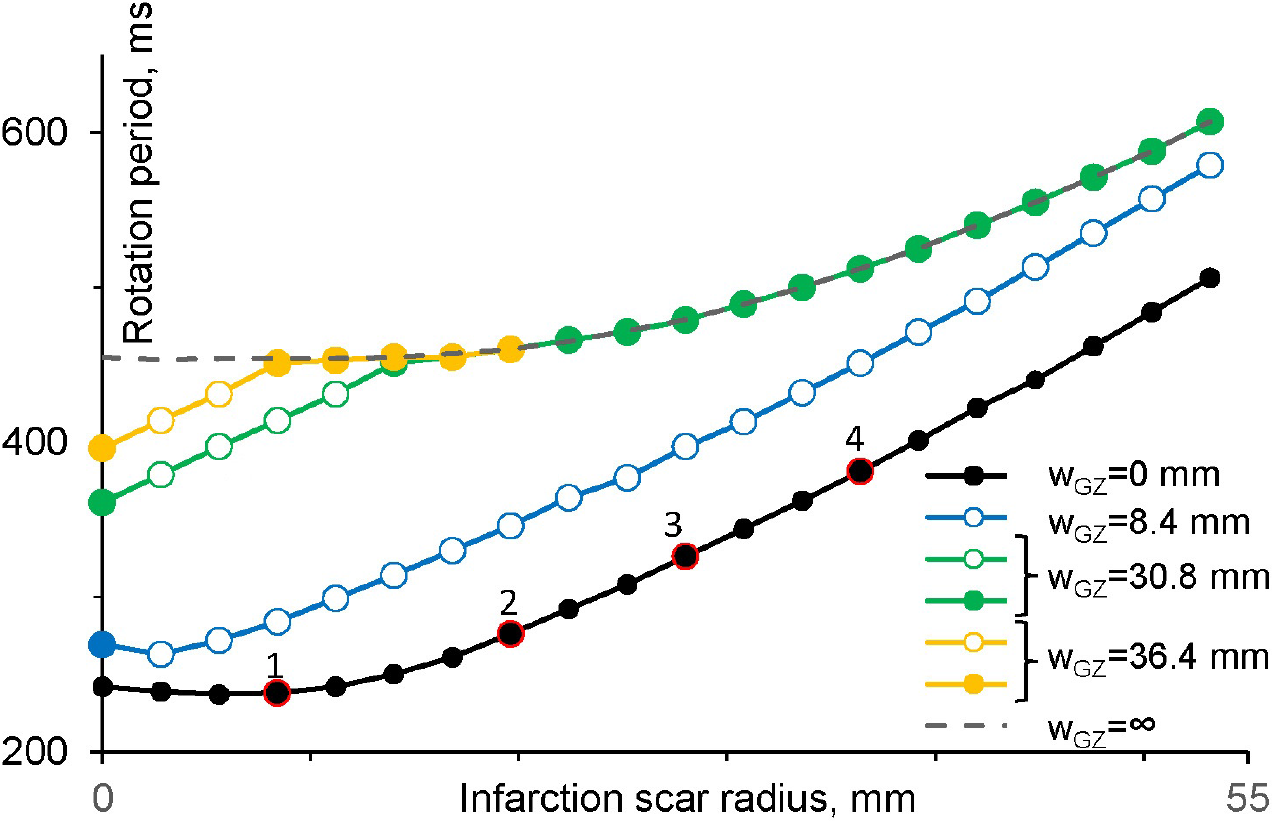
Dependence of the wave rotation period on the radius of the scar with the gray zone of different width. The black line shows cases in absence of gray zone. The gray dashed line represents periods of waves rotating around a scar in tissue with properties of the gray zone (*w_GZ_* = ∞). Filled dots (•) represent cases of the *scar rotation* regime. Open dots (◦) represent cases of the *gray zone rotation* regime. Marked dots (1)-(4) on this figure and Fig. 4 represent the same cases.

**Figure 4.**
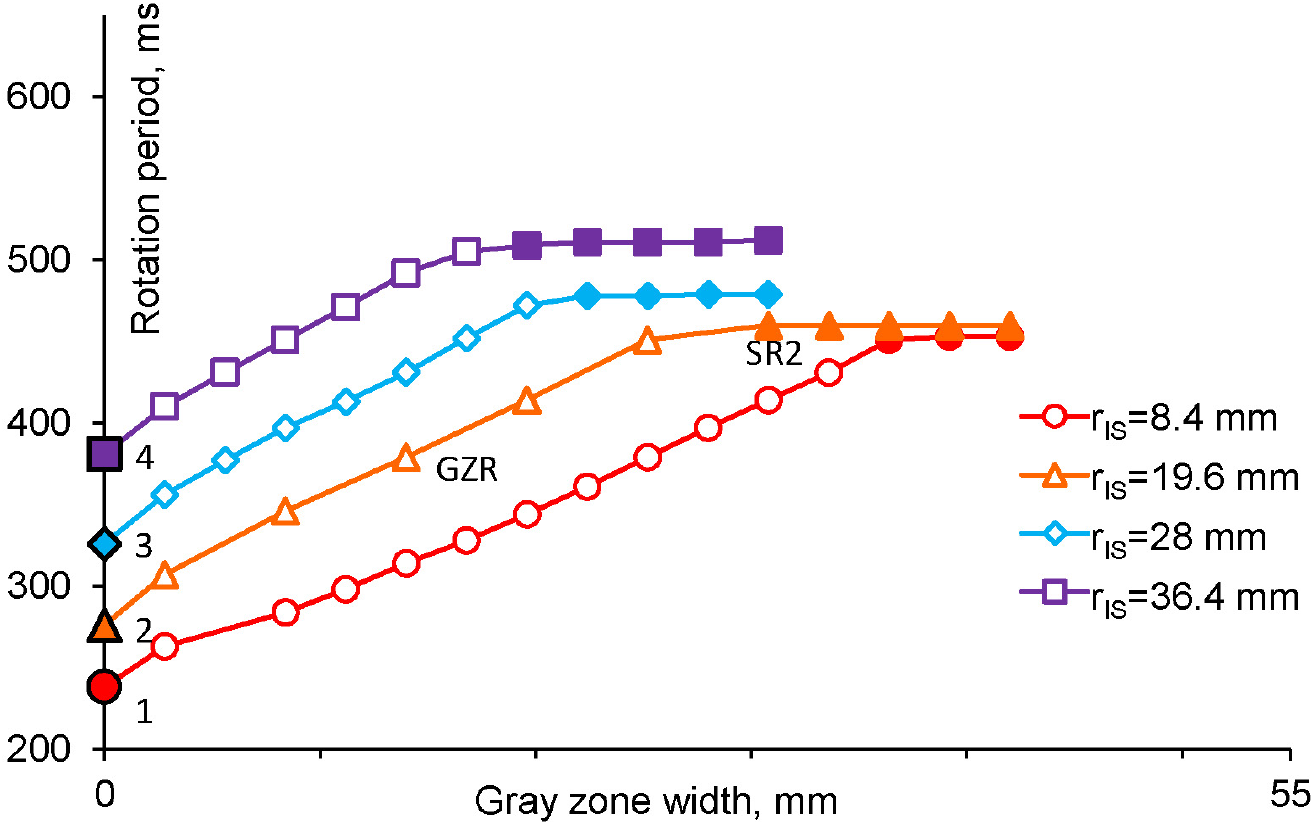
Dependence of the wave rotation period on the width of the gray zone (*w_GZ_*) in experiments with different radius of infarction scars. Red line shows case with smaller radius of infarction scar (*r_IS_* = 8.4 mm), purple line - case with bigger infarction scar (*r_IS_* = 36.4 mm). Open symbols represent cases of the *gray zone rotation* regime. Filled symbols represent cases of *scar rotation* regime. Dots (1)-(4) on this figure and on Fig. 3 represent the same cases.

Let us consider Figure 3 first. The black line shows the rotation period vs *r_IS_* in the absence of the gray zone (*w_GZ_* = 0). We see that for small *r_IS_* (*r_IS_* < 14 mm) the period is almost constant and does not depend on the size of the scar, and then dependence becomes linear with a slope of 6.95. Similarly, in the absence of normal tissue, the dashed line, (*w_GZ_* = ∞), we have a dependency with the region of almost constant period and then monotonically increasing dependency approaching linear with a slope of 5.45. (Here entire tissue has the properties of the gray zone) We will use this line later to explain the mechanisms of transitions between different wave rotation regimes. For the gray zone of width *w_GZ_* = 8.4 mm (the blue line) we have a similar dependency, with a the linear slope=6.32, however the transition to the linear part here starts for smaller infarct radius values of *r_IS_* > 2.8 mm. For the gray zone with *w_GZ_* = 30.8 and 36.4 mm (green and yellow lines, respectively), both lines approach the asymptotic dashed line at large scars. For smaller scar radius, the dependencies are linear before the green and yellow lines approach the asymptotic dashed line. Their slopes here are 6.35 and 6.5, respectively.

Now let us consider the dependence of the wave rotation period on the width of the gray zone at constant *r_IS_* (Fig. 4). For all *r_IS_*, we have similar dependencies, which at small *w_GZ_* are almost linearly increasing functions, that for larger *w_GZ_* instantly switch to the almost horizontal straight lines. We also see that for larger scars the switch to the plateau occurs at lower *w_GZ_*. In particular, for a small scar of *r_IS_* = 8.4 mm plateau phase starts around *w_GZ_* = 36.0 mm, while for the largest scar of *r_IS_* = 36.4 mm, the plateau is approached at around *w_GZ_* = 16.8 mm.

### 3.3. Period and regimes of rotation

Now let us explain the observed dependencies and relate them to the rotation regimes defined in section 3.1. Let us first consider the orange line in Fig. 4. Here the line starts at *w_GZ_* = 0 with the *scar rotation* regime and period of 276 ms. We illustrate this wave pattern in Fig. 2 (B1). When *w_GZ_* increases the wave detaches from the obstacle and starts rotating around the gray zone in the *gray zone rotation* regime. We illustrate this wave pattern with the period of rotation of 379 ms in Fig. 2 (B2). The reason for onset of such regime is the following. It turns out that for wave it is faster to rotate around the gray zone in the normal tissue, than around a scar in the gray zone tissue, as the velocity of the wave there is smaller due to longer action potential in the gray zone. Indeed, we found that the period of rotation inside the gray zone for scar of this size is 460 ms. We can see it from the dashed line in Fig.3 for *r_IS_* = 19.6 mm. This period is longer than the period of 379 ms for rotation around the gray zone, and thus wave chooses to rotate around the gray zone.

We can also explain the transition from the gray zone to scar rotation regime if we further increase the size of the gray zone. Now the period of rotation around the gray zone increases with its size, and for the gray zone of *w_GZ_* = 30.8 mm it reaches a value of 460 ms (see dot marked *SR*2 in Fig. 4). For further increase in *w_GZ_*, it will be faster for the wave to rotate around the scar than around the gray zone, and we observe the *scar rotation2* regime with period of 460 ms (Fig. 2, B3). Thus further increase in *w_GZ_* will not change the wave rotation regime and the period will remain *T* = 460 ms, as we see in Fig. 4.

Note, that as both the gray zone and scar rotations occur in the normal tissue, these regimes are closely related to each other. For example, in our case of *r_IS_* = 19.6 mm, and *w_GZ_* = 14 mm the radius of the gray zone is *r_IS_* + *w_GZ_* = 33.6 mm. From Fig. 3 we find that for scar of *r_IS_* = 33.6 mm the period of rotation is 362 ms, which is indeed close to observed value of 379 ms.

Now let consider dependencies in Fig. 3. For *w_GZ_* = 0 (the black line), the linear part of the curve is associated with the *scar rotation* regime and increasing of period of the wave is caused by increasing scar radius *r_IS_*. However, the flat part of the dependency for small *r_IS_* is associated with the *functional rotation* regime. Indeed, for small *r_IS_* the rotation effectively occurs around a circle with the radius larger than the size of the scar and thus size of the scar does not affect the rotation period An example of such effective rotation circle is shown in Fig.2 A1 by shaded blue. Similar wave patterns of the functional and scar rotation regimes are associated with the dashed curve (*w_GZ_* = ∞). However, here we obviously have the *scar rotation2* regime for large scars and the *functional rotation* regimes for small scars as the wave here propagates in the tissue with the properties of the gray zone.

For the gray zone with a large width of *w_GZ_* = 30.8 mm (see green line in Fig. 3) and large scars *r_IS_* > = 14, the curve coincides with the dashed line corresponding to the *scar rotation2* regime. However for *r_IS_* < 14 mm we have the *gray zone rotation* regime as the rotation around gray zone is faster than *scar rotation 2*. For the gray zone with the largest width of *w_GZ_* = 36.4 (yellow line in Fig. 3) we have *scar rotation 2* except smaller scars of *r_IS_* < 8.4 mm. Here we have again the *gray zone rotation*. Compared to the green curve the *scar rotation 2* here occurs at the smaller scar radius. This is also logical, as for larger *w_GZ_* the corresponding curve is closer to its asymptotic dashed curve.

Finally, let us consider the factors which determine the slopes of the dependencies of period of rotation on the scar geometry parameters.

### 3.4. Factors affecting period of rotation

To determine the factors responsible for the observed dependency of the period on the size of the scar and gray zone let us consider the following simple equation. If we assume that the velocity of wave (*v*) at the boundary of the obstacle is constant and the perimeter of the obstacle is *L*, then the period *T* is given by: *T* = *L*/*v*, or *L* = *T * v*. In turn, the velocity of the wave depends on the period of excitation *T* (conduction velocity (CV) restitution) and the curvature of the wave front [25,26]. If we neglect the dependency of the velocity on the curvature we get: *L* = *T * v*(*T*). Thus the slope of the dependency of the period on the obstacle size (shown in Fig.3) can be found from the following simple calculation:

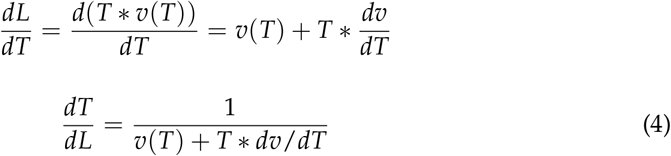

where *dv*/*dT* is the slope of the CV restitution curve.

Thus we see that the slope of the dependency of wave period *T* on the obstacle perimeter *L* is determined not only by the velocity *v* itself but also by the slope of the CV restitution curve.

We computed CV restitution curves for the normal tissue and tissue with properties of the gray zone (Fig. 5). We see that both curves are monotonically increasing functions *v*(*T*) approaching some saturated values (*v_NT_* for healthy tissue, and *v_GZ_* for gray zone tissue). Thus for large pacing periods, *dv*/*dT* approaches zero value, and according to the formulae (4) the slope *dT*/*dL* would approach the value 1/*v_NT_* or 1/*v_GZ_*, respectively. However, such saturation occurs for rather long pacing periods, which are mostly outside of the physiologically relevant values. In our case, the rotation periods are in the range of 250 < *T* < 600, where *v* strongly depends on *T* for the both CV restitution curves. Let us estimate, which terms in Eq. (4) primarily determine the slope of the dependencies in Fig.3, and explain why the curves with larger gray zone have smaller slopes than the curve without the gray zone. Let us use period *T* = 500, which is inside the region of linear dependency in Fig.3 for both curves: with and without the gray zone. For the case of a scar without the gray zone (the black curve), at *T* = 500 ms we find that *v* = 0.678, and *dv*/*dT* = 0.0003 derived from the black curve in Fig.5. We use it as in this case the wave rotates around the scar surrounded by the normal tissue. Thus in Eq. (4) the second term *T * dv*/*dT* = 0.15 is substantially smaller than the first term *v*. For a large gray zone (green curve in Fig.3) the rotation at *T* = 500 ms occurs inside the gray zone, thus we need to use the CV restitution curve for the border zone tissue (the gray curve in Fig.5). Similarly we find that for *T* = 500 ms *v* = 0.46, and *dv*/*dT* = 0.00195, thus in Eq.(4) *T * dv*/*dT* = 0.979 is higher than *v*. Thus in this case we see that now the second term *T * dv*/*dT* determines the slope of the dependency. Thus we can conclude that the lower slope of the dependency of T on L in the presence of the gray zone is a result of CV restitution slope (dv/dT), where larger derivative occurs due to ‘shift’ of CV restitution curve to the longer periods because of tissue remodelling in the gray zone (Fig.5). Note however, that linear dependencies which we see in Fig.3 for large radii actually just a coincidence for specific values of parameters. The slopes will further change and for very large values of *r* they approach the values of 1/*v_NT_* and 1/*v_GZ_* correspondingly. At that stage the slope of the dashed line in Fig.3 will be larger than that of the black line. However, it occurs far beyond the physiologically relevant values for spiral waves.

**Figure 5.**
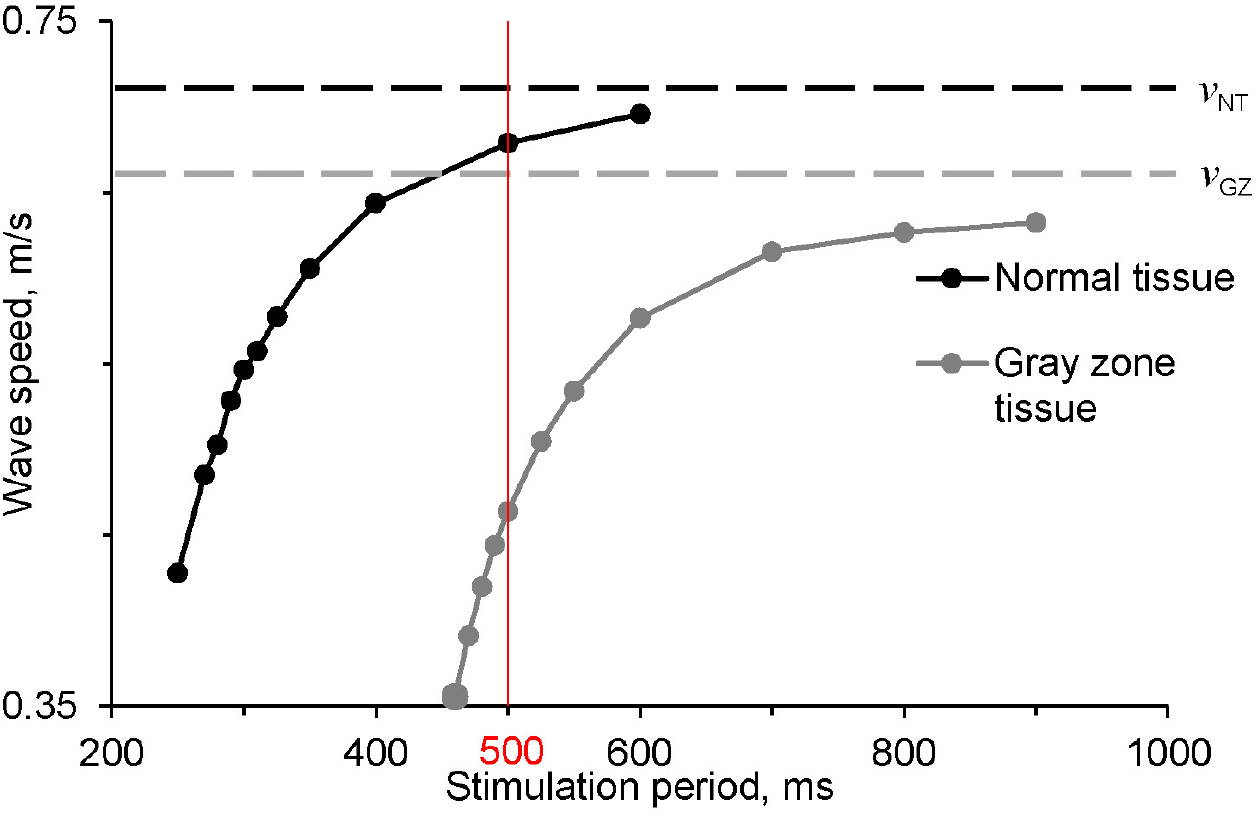
Dependence of the wave speed on the activation period of the wave (restitution curve). Wave speed in normal tissue (black) and tissue with properties of the gray zone (gray) are shown. Black and gray dashed lines show maximum speed of waves in normal tissue (*v_NT_* = 0.71 m/s) and in gray zone tissue (*v_GZ_* = 0.66 m/s) respectively.

### 3.5. Additional rotation regimes

In addition to the regimes discussed above, for few parameter values we also observed more complex rotation regimes. All these regimes occur for small scars and can be distinguished into two types: active and passive. Fig.6 A shows an example of passive regime, where we periodically see breakthrough of excitation into the gray zone, however, they do not disturb the leading edge wave rotation. Fig.6 B shows an example of active regime. Here, the wave comes outside the gray zone, actively interacts with the leading edge of excitation and changes the trajectory of the wave rotation. As a result we observe a non-monotonical dependency of the period on the size of the scar. However, the active regime was not frequently observed, we saw it only in 3 cases (out of 360) with perimeter of gray zone 52.7 mm (*r_IS_* = 0 mm, *w_GZ_* = 8.4 mm; *r_IS_* = 2.8 mm, *w_GZ_* = 5.6 mm; *r_IS_* = 5.6 mm, *w_GZ_* = 2.8 mm), thus they may have limited importance for overall dynamics.

**Figure 6.**
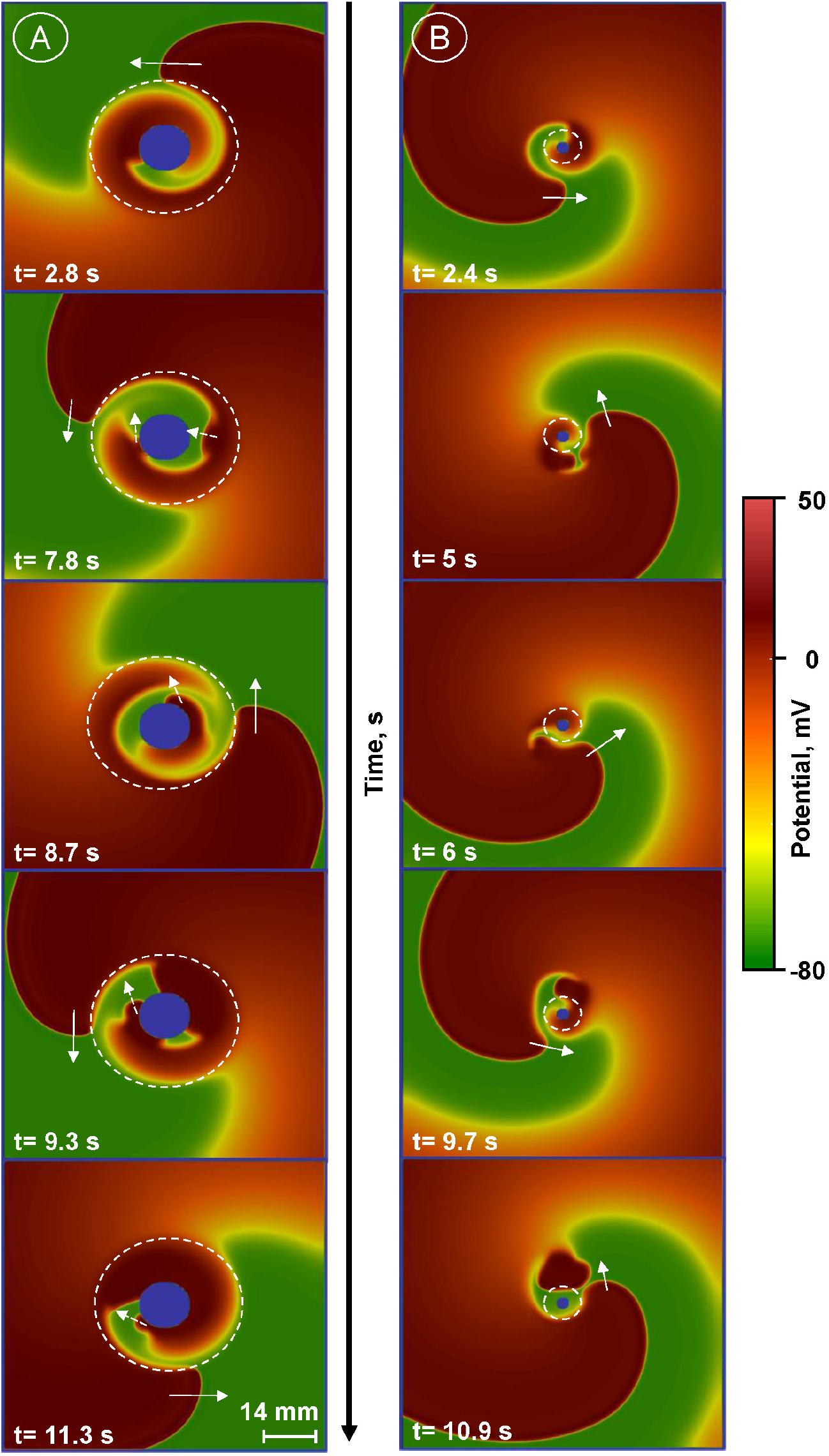
Examples of waves with dynamical instabilities. Border of the gray zone is depicted by the dashed line. Arrows show the direction of the wave rotation. Dashed arrows show wave direction inside the gray zone. A: a passive regime. The breakthrough waves marked by dashed arrows do not exit the gray zone (*r_IS_* = 11.2 mm, *w_GZ_* = 25.2 mm). B: an active regime. The breakthrough waves exit the gray zone and force the wave rotate at some distance from the border of the gray zone (*r_IS_* = 2.8 mm, *w_GZ_* = 5.6 mm.) An illustration of such dynamics can be found in the Supplement movies (movies S3 and S4).

## 4. Discussion

In this paper we perform a detailed study of wave dynamics rotating around a region, which generically represents a post-infarction scar and includes a compact inexcitable obstacle surrounded by a gray zone with modified cellular activity as compared with the rest of normal tissue. We found that all observed dynamics can be subdivided into several main classes. These classes naturally occur if we combine properties of wave rotating around inexcitable obstacle and effects of tissue heterogeneity on the wave propagation.

In the absence of tissue heterogeneity, the period of wave rotating around an obstacle is mainly determined by the velocity of the wave and the perimeter of the obstacle [10]. This is a *scar rotation* regime, which was observed in previous studies. However, compared to earlier studies [10] we show that one needs to take into account the dependency of the velocity on the period (CV restitution), which substantially affects the period of wave rotation. Another observed regime of *functional rotation*, was found in early studies in the reaction diffusion models [11]. It was shown that for small obstacles the period of rotation does not decrease anymore, wave detouches from the obstacle and rotates around a circle which exceeds the obstacle size. In [11] it was also shown that detachment from the obstacle is associated with a large increase in the period of rotation. We did not observe such period increase in our study. This probably occurs because in [11] a two variable model with lower excitability was used. In our study we used an ionic model with high excitability, and thus boundary effects of the obstacle on the wave propagation velocity in our case are less pronounced.

In the presence of the gray zone, the situation is more complex and, as far as we aware, was not yet analyzed in details. Our study shows that here we can observe additional rotation regimes, which we call *gray zone rotation* and *scar rotation 2* regimes. We show that dependencies of period on the size of the scar and gray zone are bounded by two asymptotic solutions for homogeneous tissue: rotation of wave around a scar in normal tissue and rotation of wave around a scar in tissue with properties of the gray zone. As we show in Fig. 3 all curves for different sizes of the gray zone are located between these two asymptotic lines and the slopes of these lines bound from above and below the slopes of all other curves. In addition, we found a new regime, *gray zone rotation*. In that case, the wave rotates as if the compact scar coincides with the gray zone. In the *gray zone rotation* regime the period of rotation depends on the perimeter of the gray zone, and not on the size of the scar. We illustrate it in Fig. 7, where we show the dependence of the period on the perimeter of the gray zone for various sizes of the scar. We see that all curves in certain range of parameters almost coincide, which one more time confirms that here the perimeter of the gray zone, and not the scar size determines the period of rotation. However, we also see that this curves with the gray zone are little above the dashed line, representing the case of pure scar rotation. It indicates that the leading edge of rotation for the *gray zone rotation* regime is a little outsize the boundary of the gray zone, which maybe a result of the electrotonic effects [27].

**Figure 7.**
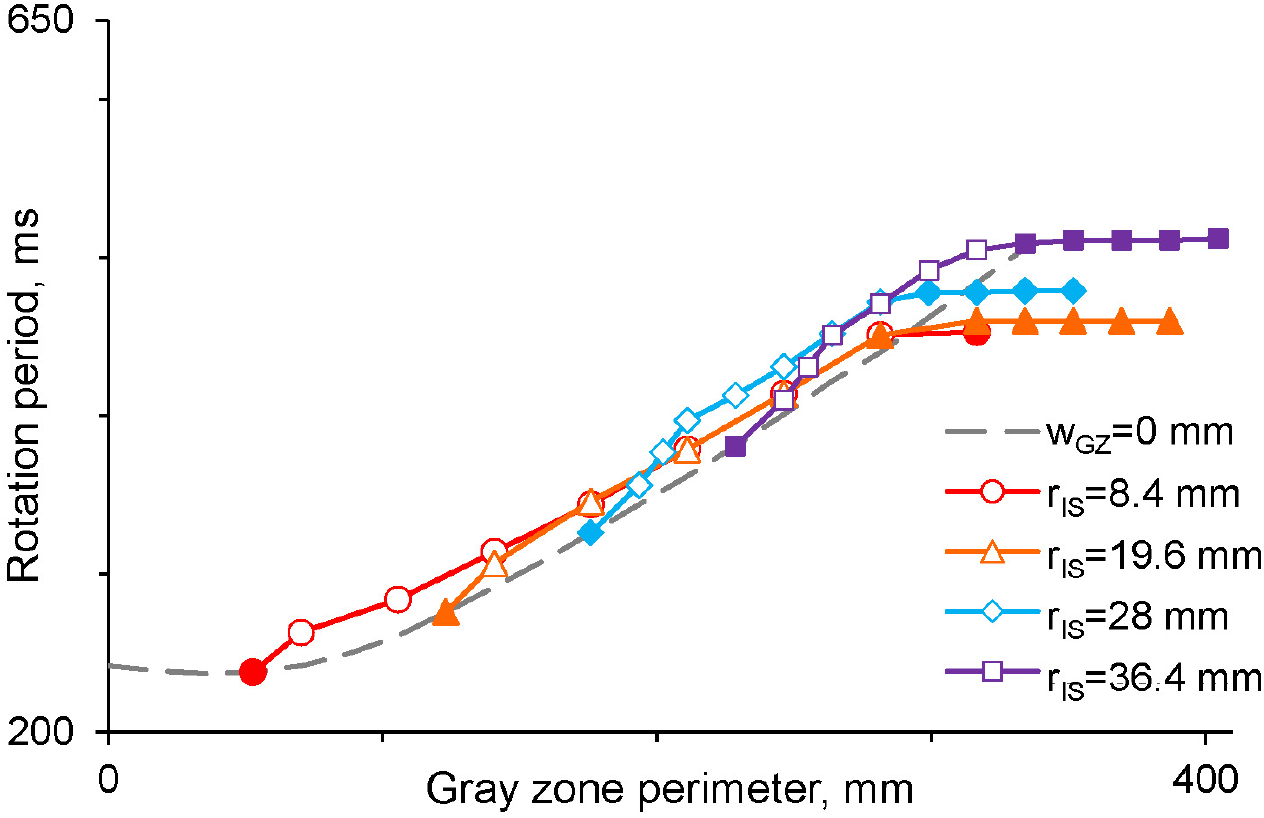
Dependence of the wave rotation period on the perimeter of the gray zone for different radii of infarction scar. Red line shows case with smallest radius of scar core, purple line - case with biggest scar core. The dashed line connects points where the gray zone is absent (*w_GZ_* = 0 mm).

Important rule for transition between the gray zone and scar rotation regimes is the minimal period principle. In fact, if rotation in normal tissue along the gray zone is faster than the rotation around the scar in the gray zone, then the *gray zone* regime will occur. Such transition is clearly seen in Fig.3 and Fig. 4 where we see change from one type of dependency to another one. Note, this also implies that transitions between the regimes strongly depend on the relation of the velocity of the wave inside and outside the gray zone. Here we considered only one type of cellular remodeling based on the data reported in [21,22], where action potentials are longer in the gray zone. However in [28] it was shows that remodelling in the gray zone can also result in different changes in the properties of cardiac tissue and opposite effects on action potential duration. Thus it would be interesting to study similar regimes of wave rotation for different properties of cardiac tissue in the gray zone. In addition, the gray zone can also include fibrosis and reduced number of gap junctions. It also can decrease the propagation velocity and thus make *gray zone rotation* regime more essential.

The dynamics of rotating waves around an obstacle was studied in [14], where a general theory for the dynamics and instability of pinned spirals was developed. In particular, it was shown that the destabilization is better understood by the implementation of a mapping rule and dimension reduction.

Also transition between scar and functional rotation when the radius of the obstacle decreases were studied in [29]. There a sequence of transitions, from periodic motion to a modulated period-2 rhythm, and then to spiral wave breakup [30,31] were observed. It would be interesting to study how such dynamical instabilities will manifest itself in the presence of tissue heterogeneity around a scar, as considered in our paper.

It would be also interesting to study such rotational activity in anatomical models of the heart [22,32] and also include additional types of regional cellular heterogenety in the normal tissue as considered in [33,34]. Note that cellular heterogeneity, especially involving pathological elongation of the action potential, can lead to early afterdepolarizations and new types of spatio-temporal dynamics [35].

In conclusion, we performed the first detailed studies of regimes of wave rotating around an obstacle surrounded by heterogeneous ventricular myocardial tissue, which is a generic model of the infarction scar with the gray zone. We found that the main regimes here include rotation around the compact scar and also rotation around the gray zone. We explained dependency of the rotation period on the size of the scar and showed that it is strongly affected by the conduction velocity restitution. We also show that the transition between the regimes can be understood and predicted from the minimal period principle.

## Supplementary Materials

The following are available at https://www.mdpi.com/2227-7390/1/1/0/s1, Video S1: Example of wave rotating in *gray zone rotation* regime. Infarction scar size:*r_IS_* = 19.6 mm. Gray zone width *w_GZ_* = 14 mm. Video S2: Example of wave rotating in *scar rotation 2* regime. Infarction scar size:*r_IS_* = 19.6 mm. Gray zone width *w_GZ_* = 42 mm. Video S3: Example of the wave with passive dynamic instability in gray zone. Infarction scar size:*r_IS_* = 11.2 mm. Gray zone width *w_GZ_* = 22.4 mm. Video S4: Example of the wave with active dynamic instability in gray zone. Infarction scar size:*r_IS_* = 2.8 mm. Gray zone width *w_GZ_* = 5.6 mm.

## Author Contributions

Conceptualization, A.P., D.M. and O.S; formal analysis, P.K., D.M.;Investigation, P.K.; methodology, A.P. and P.K.; software P.K., A.D. and D.M.; supervision, A.P. and O.S.; visualization, P.K. and A.D.; writing–original draft preparation, P.K., A.P.and O.S.; writing–review and editing, P.K., A.P. and O.S.; All authors have read and agreed to the published version of the manuscript.

## Abbreviations

Abbreviations

The following abbreviations are used in this manuscript:

CV: Conduction velocity
FR: Functional rotation
GZ: Gray zone
GZR: Gray zone rotation
IS: Post-infarction scar
NT: Normal tissue
SR: Scar zone rotation
SR2: Scar rotation 2

